# Eccentricity strongly modulates visual processing delays

**DOI:** 10.1101/2023.09.30.559991

**Authors:** Johannes Burge, Callista M. Dyer

## Abstract

The temporal dynamics of visual information processing varies with the stimulus being processed and with the retinal location that initiates the processing. Here, we present psychophysical data with sub-millisecond precision showing that increasing eccentricity decreases the delay with which stimuli are processed. We show that, even within the central +/-6º of the visual field, processing delays change by a factor of up to three times. A simple model, grounded in retinal physiology, provides a good account of the data. The relative delays are on the order of only milliseconds. But if later processing leaves the delays unresolved, they can cause dramatic misperceptions of motion and 3D layout. We discuss the implications for how the human visual system solves the temporal binding problem across eccentricity. The results highlight the severe computational challenge of obtaining accurate, temporally-unified percepts of the environment with spatiotemporally-staggered processing across the visual field.

## Introduction

Catching a ball at dusk is more difficult than at high noon. Starting with the photoreceptors, visual processing is slower when overall light-levels are low (Lit, 1949; Baylor & Hodgkin, 1973; Drum, 1984; Sinha et al., 2017; Burge et al., 2019). When sensors signal the rest of the nervous system with more delay, there is less time to react. The speed of visual processing depends on many stimulus properties other than overall light-level. Different contrasts (Nachmias, 1967; Reynaud & Hess, 2017), colors (Cottaris & DeValois, 1998), level of detail (i.e. spatial frequency) (Harwerth & Levi, 1978; Min et al., 2021; Chin & Burge, 2022), and levels of image sharpness (Burge et al., 2019; Rodriguez-Lopez et al., 2020; Rodriguez-Lopez et al., 2023) are all processed with different temporal dynamics (e.g. delays and temporal blurring). In any natural image, these stimulus properties tend to vary substantially across the visual field (Fig. 1A, top). The sky may be bright, blue, and devoid of detail; the underbrush may be dark, green, and full of detail. So neural signals carrying information about each location of a given image will tend to move through the early stages of the visual system in a temporally staggered fashion (Fig. 1A, middle). However, it is not well-understood how visual field location *itself* modulates the speed with which any given stimulus property is processed (Fig. 1A, bottom). Will a given stimulus-based discrepancy in processing speed be increased, decreased, or unaffected by where stimuli are imaged in the visual field? There are conflicting results in the literature (Rutschman, 1966; Osaka, 1976; Virsu et al., 1982; Carrasco et al., 2003).

**Figure 1.**
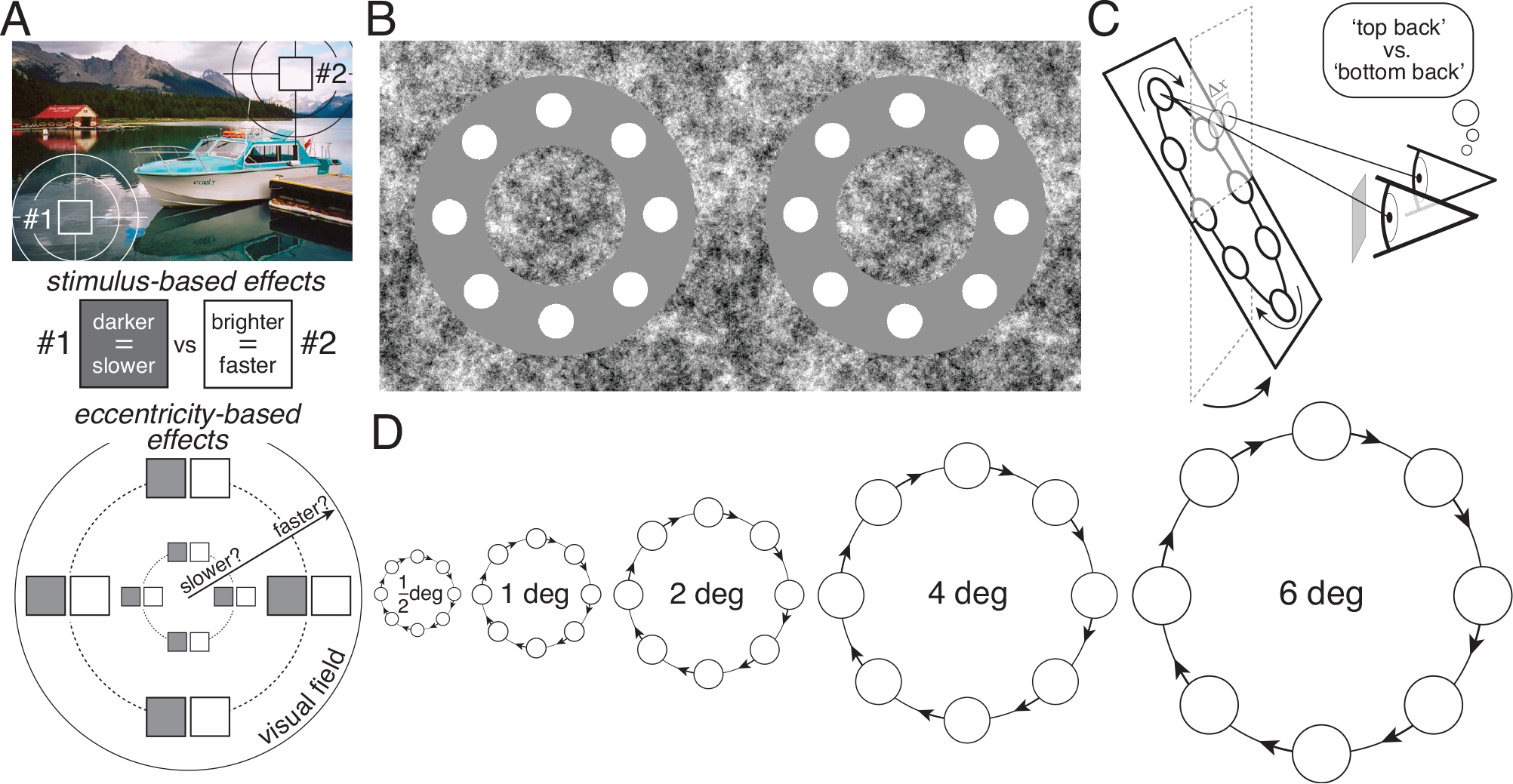
Processing speed, stimulus, and task. **A** Stimulus properties vary with location in natural scenes (top). Dark stimulus patches (#1) are processed more slowly than bright stimulus patches (#2) when fixated (middle). It is largely unknown how visual field location impacts processing speed (bottom). **B** Left- and right-eye onscreen stimuli. Free-fuse to see the stimuli in depth. Divergent-fusing will produce a ‘top-back’ percept. Cross-fusing will produce a ‘bottom-back’ percept. **C** If the left eye receives less light than the right eye, the left-eye image will be processed with more delay. With moving stimuli, a neural disparity will occur, and the beads will be perceived as rotated ‘top back’ with respect to the screen. The effective neural disparity at a given bead location Δ*x* is given by *ν*_*x*_Δ*t* the product of the horizontal velocity component of the bead motion, and the interocular neural delay assuming that onscreen delays are zero. The critical onscreen delay was that which nulled the neural delay, and made the beads appear to rotate in the plane of the screen. **D** Critical onscreen delays were measured for a five different eccentricities (0.5º, 1.0º, 2.0º, 4.0º, 6.0º). These critical onscreen delays should be equal in size but have the opposite sign of the neural delay. Circular bead sizes (10, 20, 40, 80, 120arcmin) and rotational speeds (1, 2, 4, 8, 12deg/sec) scaled exactly with eccentricity.

Here, we ask how visual eccentricity modulates the speed with which a given stimulus property is processed. To probe temporal processing in peripheral regions of the visual field, we had subjects binocularly view, with luminance differences between the eyes, a stimulus composed of a rotating set of white circular beads (Fig. 1BC). When the beads rotate in the plane of the screen, they are inaccurately perceived as rotating in a plane that is pitched back from the screen. This illusory percept is a modified version of the classic Pulfrich effect (Pulfrich, 1922; Lit, 1949; Burge et al., 2019). It occurs because the image in the darker eye is processed more slowly than that in the brighter eye. The difference in processing speed between the eyes—the neural interocular delay—causes a neural disparity for moving stimuli, which leads to the misperceptions.

In a psychophysical experiment, we use this stimulus to estimate the luminance-difference induced discrepancy in processing speed between the eyes—the neural interocular delay—and then determine whether this interocular delay changes with eccentricity. The estimates of these delays have sub-millisecond precision, enabling precise characterization of how processing speed changes with eccentricity. We find that the eccentricity at which a given stimulus property is processed strongly modulates the speed of processing (Carrasco et al., 2003). Delays decrease by approximately three times within only the central ±6º of the visual field. A simple model suggests that eccentricity-dependent changes in the light-sensing properties of the cone photoreceptors can account for these effects (see below). The absolute differences in processing speed at different eccentricities may be tiny (i.e. on the order of milliseconds), but if they are not resolved, the perceptual consequences of these spatially-variant delays can be profound.

Interestingly, because percepts tend to be accurate and temporally unified, the delays do seem to be resolved in most viewing situations. Rigidly moving scenes, for example, tend to be perceived as moving rigidly, despite the fact that the temporal dynamics of visual processing depend on the stimulus and its location in the visual field. The present findings challenge vision science to develop a rigorous understanding of how temporally staggered signals caused by the same event are synchronized and bound together in time.

The current findings also help plug a gap in the literature. The impact of eccentricity on some aspects of temporal processing has been well-characterized. Temporal sensitivity—which indicates the smallest temporal modulation that is detectable—is well-known to increase with eccentricity: sensitivity to flicker, for example, is higher in the periphery than in central vision (Hartmann et al., 1979; Kelly, 1984). (This is why some fluorescent lights in badly lit bars can be seen to flicker from the corner of one’s eye.) Temporal delay, and how it is impacted by eccentricity, is much less well understood. Few studies have examined how processing speed changes across the visual field (but see Osaka, 1976; Carrasco et al., 2003; Jovanovic & Mamassian, 2020; Upadhyayula, Phillips, Flombaum, 2023). Fewer still contain within-subjects measurements with the resolution required to precisely characterize how the speed of visual processing changes across the visual field. The current study reports just such measurements.^1^

## RESULTS AND DISCUSSION

### Psychophysical Results: Main Experiment

To measure the eccentricity-dependent delays in the human visual system, we designed a one-interval two-alternative forced-choice (2AFC) experiment. The stimulus was a rotating set of circular beads that followed the center of an annulus-shaped ‘racetrack’ (Fig. 1BC). On each trial the beads rotated either clockwise or counter-clockwise while the subject fixated a central dot. Trial duration was 250ms, the approximate length of a human fixation. A haploscope enabled dichoptic stimulus presentation. Luminance differences between the eyes were introduced programmatically. These luminance differences cause one eye’s image to be processed with more delay than the other. This neural interocular delay, in turn, causes the set of beads to be perceived as moving in a plane different than the screen plane. The task was to report, on each trial, whether the plane in which the beads were moving appeared to be pitched ‘top-back’ or ‘bottom-back’ relative to the screen (Fig. 1C). Neural delay can be compensated for by manipulating the onscreen timing of stimulus presentation. A neural delay in the left eye, for example, can be nulled by advancing the left-eye image onscreen. The aim of the experiments was to find, for each luminance-difference and eccentricity, the onscreen delay that caused the beads to be perceived as moving in the plane of the screen. These critical onscreen delays should be equal in size, and opposite in sign, to the neural delay in each condition.

On each trial of the experiment, onscreen interocular delay was manipulated in pseudo-random fashion (see Methods). If it was zero, the onscreen binocular disparities on each frame specified that the beads were rotating in the plane of the screen (see Methods). If the onscreen-delay was non-zero, onscreen binocular disparities specified that the plane in which the beads were rotating was pitched top-back or bottom-back relative to the screen plane (Fig. 1C). In each condition, we computed the proportion of trials in which subjects responded ‘top-back’ as a function of onscreen interocular delay. Data was collected for four interocular luminance differences and at each of five eccentricities. Different eccentricities were probed by changing the radius of the annulus-shaped racetrack (Fig. 1D).

The psychometric functions for the first human subject are shown in Fig. 2A. The functions get closer together as eccentricity increases, indicating that a fixed luminance difference has less effect on processing delays in the visual periphery. We summarize performance in each condition by computing the point of subjective equality (PSE). The PSE is the onscreen interocular delay corresponding to the 50% point on the psychometric function (colored arrows), and specifies the delay that is required to null the neural disparity such that the beads are perceived as rotating in the plane of the screen. (Note also that in addition to getting closer together, the psychometric functions get steeper with eccentricity. Delay discrimination thresholds are thus smaller in the periphery, a finding consistent with there being better temporal contrast sensitivity (e.g. higher flicker fusion frequencies) in the visual periphery (Kelly, 1984); see Supplement Fig. S1).

**Figure 2.**
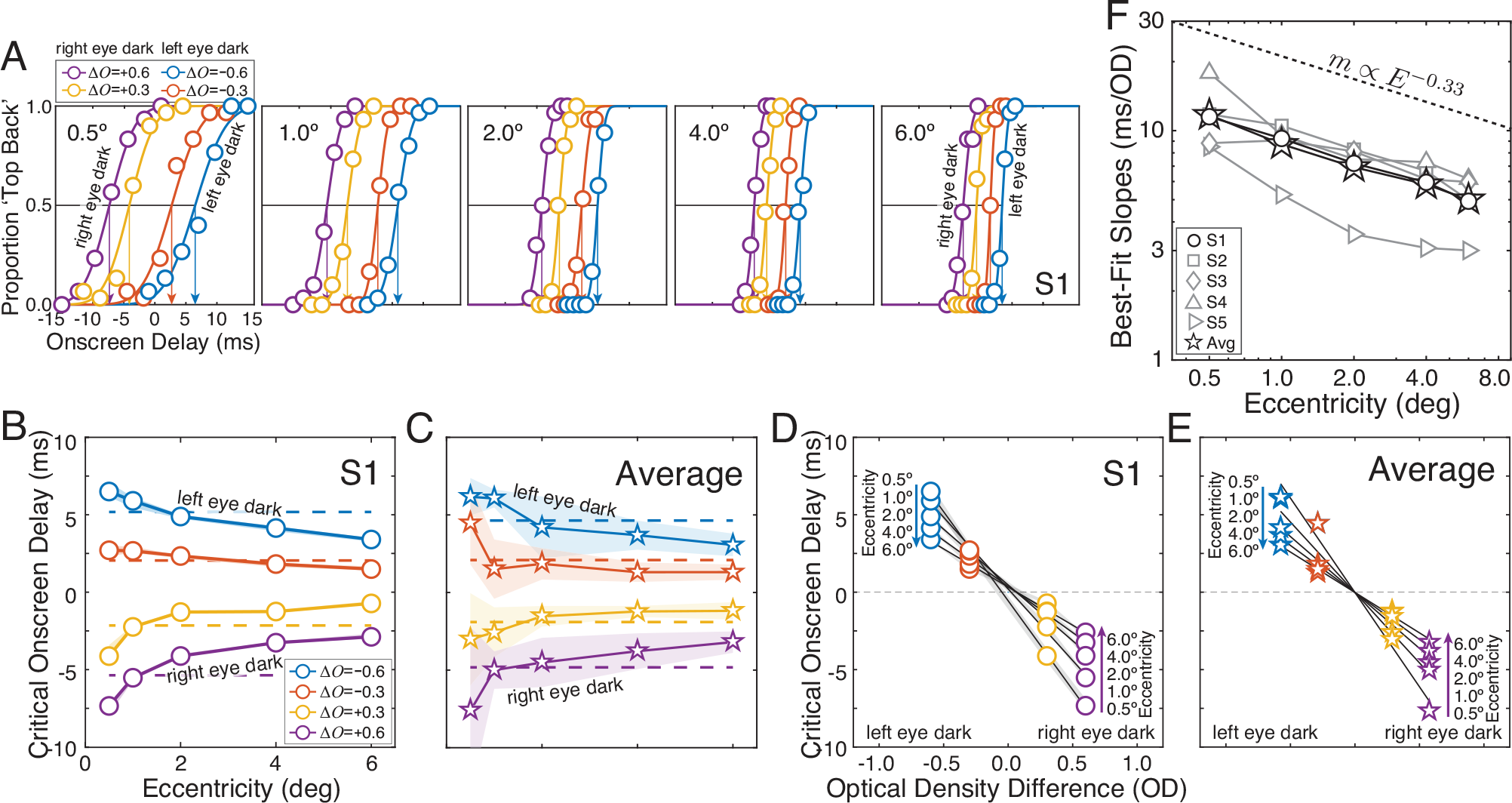
Main experiment. **A** Psychometric functions for the first human subject. Proportion ‘top back’ chosen as a function of onscreen delay for each luminance difference between the eyes (colors) and each eccentricity (subplots) for subject S1. Luminance differences are expressed as interocular differences in optical density (Δ*O*=[-0.6,-0.3,0.3,0.6], colors; see Methods). Points of subjective equalities (PSEs; arrows) indicate the critical onscreen delays required to make the stimulus appear to rotate in the plane of the screen. **B** Critical onscreen delay as a function of eccentricity for the four interocular luminance differences. Delays decrease systematically with eccentricity. Shaded regions indicate bootstrapped 68% confidence intervals (approximately ±1 standard error). In cases where shaded regions are not visible, it is because the confidence interval is smaller than the datapoint. **C** As in B, but for averaged data across subjects. Shaded regions indicate standard deviations across subjects. **D** Interocular delays as a function of interocular difference in optical density, for five different eccentricities (replotted data in A). Best-fit regression lines, computed via weighted linear regression, are also shown. The slopes of the best-fit lines decrease systematically with eccentricity, again indicating that the same luminance difference causes smaller interocular delays as eccentricity increases. **E** As in D, for group averaged data. **F** Best-fit slopes (see D and E) at each eccentricity for all individual subjects and the average data (symbols and solid lines). The best-fit power function—computed from the average data (stars)—is also shown (dashed black line; see Methods). For the group average data, the delays caused by a given luminance difference decrease with eccentricity raised to a power of -0.33. (Individual subject best-fit powers, in order, are *m* = [-0.34, - 0.34, -0.21, -0.34, -0.40].) Hence, 8-fold increases in eccentricity are associated with 2-fold decreases in visual processing delay.

Critical onscreen delays (i.e. PSEs) decrease systematically with eccentricity (Fig. 2B). For each of four different luminance differences (colors), as eccentricity increases from 0.5º to 6.0º (panels), the interocular delay decreases by ∼2.5x. For example, when the right eye received 75% less light than the left (*ΔO=*+0.6) the amount by which the left-eye image had to be delayed onscreen decreased from 7.3ms near the fovea (0.5º of eccentricity) to only 2.9ms in the near periphery (6.0º of eccentricity). The group averaged data follows a similar pattern (Fig. 2C). (Data from each of the five individual subjects also show a similar pattern; see Supplementary Fig. S2.) Replotting the data from Fig. 2BC reveals that the critical onscreen delays change linearly with optical density difference (*ΔO*), and that the best-fit regression lines (see Methods) have slopes that decrease systematically with eccentricity (Fig. 2DE). (This decrease in the slopes is reflected in the psychometric functions by how they get closer together as eccentricity increases; see Fig. 2A.)

To summarize the influence of eccentricity on visual processing delays, we plot how the slopes of the best-fit lines (see Fig. 2DE) change with eccentricity for the group average and each of the five human subjects (Fig. 2F). The proportional (and absolute) changes are largely similar across individuals. The data is (approximately) a straight line on a log-log plot, and hence is well described by a power function *m*(*E*) ∝ *E*^*p*^ over the range of tested eccentricities. A fixed proportional change in eccentricity thus causes a fixed proportional change in processing delay. A power of *p*=-0.34 provides the best fit to the first subject’s data (see Fig. 2D). A power of *p*=-0.33 provides the best fit to the group averaged data (see Fig. 2E). This best-fitting power entails that an 8-fold increase in eccentricity entails a 2-fold decrease in visual processing delay. The data strongly suggest that eccentricity lawfully changes the speed at which a given stimulus property is processed.

### Psychophysical Results: Size-Control Experiment

Well-designed scientific studies attempt to rule out obvious potential confounding factors as alternative explanations of the results (Burge & Burge, 2022). The stimuli in the main experiment were composed of beads with sizes that covaried (i.e. scaled perfectly) with the eccentricity being probed (see Fig. 1D). But stimulus-size itself has been identified as a factor that can modulate the temporal properties of visual processing, both in psychophysics and neurophysiology (Virsu et al., 1982; Watson, 1986; Carrasco et al., 2003; Dacey et al., 2000; Bair, Cavanaugh, Movshon, 2003; Churan, Richard, Pack, 2009). To justifiably conclude that eccentricity is responsible for the effects reported here, bead-size must be controlled for.

We conducted a size-control experiment that was identical to the original experiment, except that bead-size was held constant at all eccentricities (Fig. 3A). If the data from the control experiment are the same as those from the original experiment, the change in bead-size with eccentricity can be ruled out as an explanation of the results. Data from the size-control and original experiments are shown in Fig. 3B (left and right subplots, respectively). The patterns are similar: the changes in critical onscreen delay from 1.0º to 6.0º of eccentricity are nearly identical in the size-control and original experiments.

**Figure 3.**
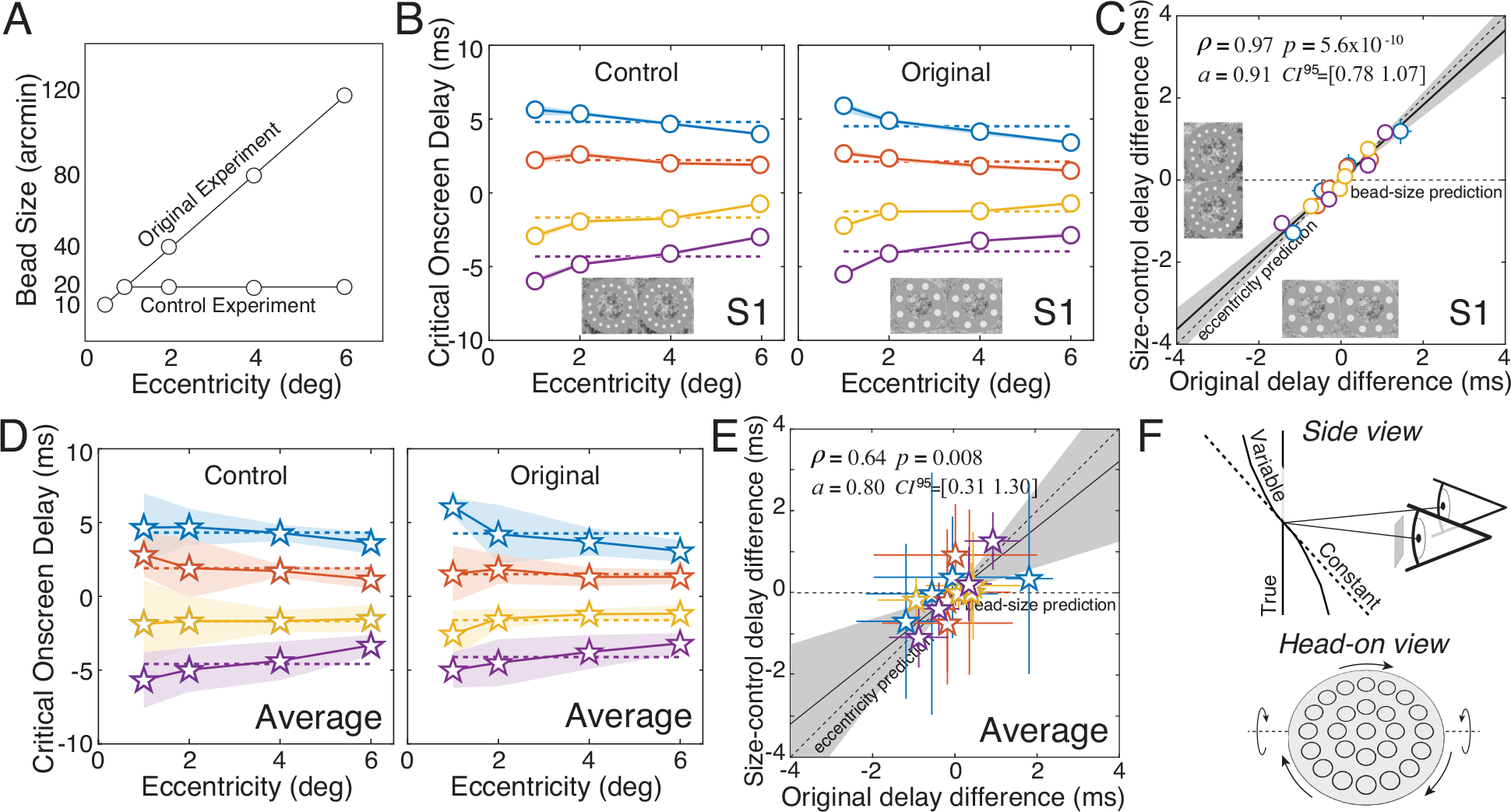
Size-control experiment. **A** In the original experiment, bead size increased in proportion to eccentricity. In the control experiment, bead size was held constant at all eccentricities. **B** Critical onscreen delays as a function of eccentricity at each luminance difference (colors) in both the control experiment (left) and the original experiment (right) for the first human subject. **C** Delays from the size-control experiment plotted against delays from the original experiment, after having regressed out average delays due to luminance differences (dashed lines in B). The eccentricity-dependent changes in critical onscreen delay were significantly correlated between the two experiments (*ρ*=0.97, *p*=5.6×10^−10^). The slope *a* of best-fit regression line, obtained using Deming regression, was not significantly different from 1.0 (*a* =0.91, *CI*^*95*^=[0.78 1.07]). Shaded regions show 95% confidence intervals on the best-fit slopes obtained from 1000 bootstrapped datasets. **D** As in B but with group averaged data. Shaded regions indicate the standard deviation across subjects in each condition. **E** As in C, but for group averaged data (*ρ*=0.64, *p*=0.008; *a* =0.80, *CI*^*95*^=[0.31 1.30]). The individual subject data, in order, are best-fit by *a* =[0.91, 1.10, 1.49, 0.96, 3.25] (see Supplement Fig. S2D). The evidence indicates that bead-size has no significant effect on delay. **F** Predicted 3D percepts of a binocularly viewed spinning object (true), assuming interocular delays that either are constant with (dashed) or vary with (solid) eccentricity. All predictions assume that no computations subsequent to the delays function to eliminate them.

To more rigorously compare the data, we subtracted off the effect of luminance differences (Fig. 3B, dashed lines), and then plotted how critical onscreen delay changed with eccentricity in the two experiments against each other. We then computed the correlation and best-fit regression line using Deming regression (Fig. 3C), which is appropriate when both variables are uncertain due to measurement error, as they are here. (Failing to account for such measurement error will lead to systematic underestimation of the slope of the best-fit regression line relating the true values of the variables. Failing to regress out the effect of luminance differences would artificially inflate the correlation and increase the slope the best-fit regression line.)

For the first human subject, the data from the two experiments are nearly identical. The correlation between eccentricity-based delay changes was clear and significant (*ρ*=0.97, *p*=5.6×10^−10^). Perhaps more importantly, the best-fit regression line was not significantly different from the unity line (*a*=0.91, *CI*^*95*^=[0.78 1.07]), which is what one would expect if bead-size had no influence on the results. Similar patterns of results are obtained for the group averaged data (Fig. 3DE) and individual subject data (see Supplementary Fig. S2). The control experiment shows that, in this task, bead-size changes do not alter visual processing delays. Bead-size, therefore, is not confounding the results. (Carrasco et al. (2003) reports that stimulus size does impact processing delays. Although there are differences in the task (target detection), data-collection paradigm (response-times), analyses (speed-accuracy-based), and the potential for attention-based effects, we do not have an explanation for the discrepant results; also see Footnote #1.)

What are the consequences of these eccentricity-dependent processing delays for visual perception? Consider a pepperoni pizza that has been tossed into the air and is spinning on its axis. If subsequent processing leaves eccentricity-dependent delays unresolved, striking perceptual errors will result. For the situation depicted in Fig. 3F, the flat pizza will be seen to have an S-shape and an average 3D orientation very different from the true orientation, or from what would be predicted with interocular delays that are constant across eccentricity. But this particular perceptual illusion, and others that are predicted from spatially-variant temporal processing, do not typically occur in normal viewing situations. For temporally staggered signals from across the visual field to contribute to a coherent unified percept they must, at some point in visual processing, be bound together in time. An important direction for future research is to develop an understanding of how these temporal-binding mechanisms work, and under what circumstances they fail.

### Retinal-physiology-based model

Why does eccentricity impact visual processing delays? More specifically, why does a given luminance difference induce larger delays in the central than in the peripheral visual field? We now ask whether the manner in which basic properties of the retina change with eccentricity can account for the processing delays that we have measured psychophysically.

We pursue this line of inquiry because the classic Pulfrich effect is widely thought to have its origins in the processing of the retina itself (Prestrude & Baker, 1968; Prestrude, 1971; Rogers & Anstis, 1972; Mansfield & Daugman, 1978; Lennie, 1981; Bolz, Rosner, Wassle, 1982; Wolpert et al., 1993). This view is based primarily on evidence that changes to retinal response and to the Pulfrich effect follow similar time-courses when subjected to conditions prompting short- and long-term adaptation. However, the Pulfrich effect has not previously been measured in the visual periphery. So, although a great deal is known about how various aspects of retinal physiology change with eccentricity, it is not known whether the effects reported here are accounted for by how the temporal properties of response change across the retinal surface.

To investigate, we built a simple model. The model incorporates how three major facets of retinal physiology change with eccentricity: photoreceptor inner-segment diameters increase, photoreceptor outer-segment lengths decrease, and macular pigment density decreases (Curcio et al., 1990; Banks et al., 1991; Putnam & Bland, 2014; Cottaris et al., 2019) (Fig. 4A). Consistent with measurements of cone phototransduction (Sinha, 2022; Baudin, Angueyra, Sinha, Rieke, 2019), the model also assumes that, for a given light-adapted state at a moderate photopic light-level in line with our experimental conditions (∼100cd/m^2^), i) each photoreceptor must catch a threshold number of photons before a response is elicited, and ii) that this threshold number of photons does not change within the central ±20º of the visual field.

**Figure 4.**
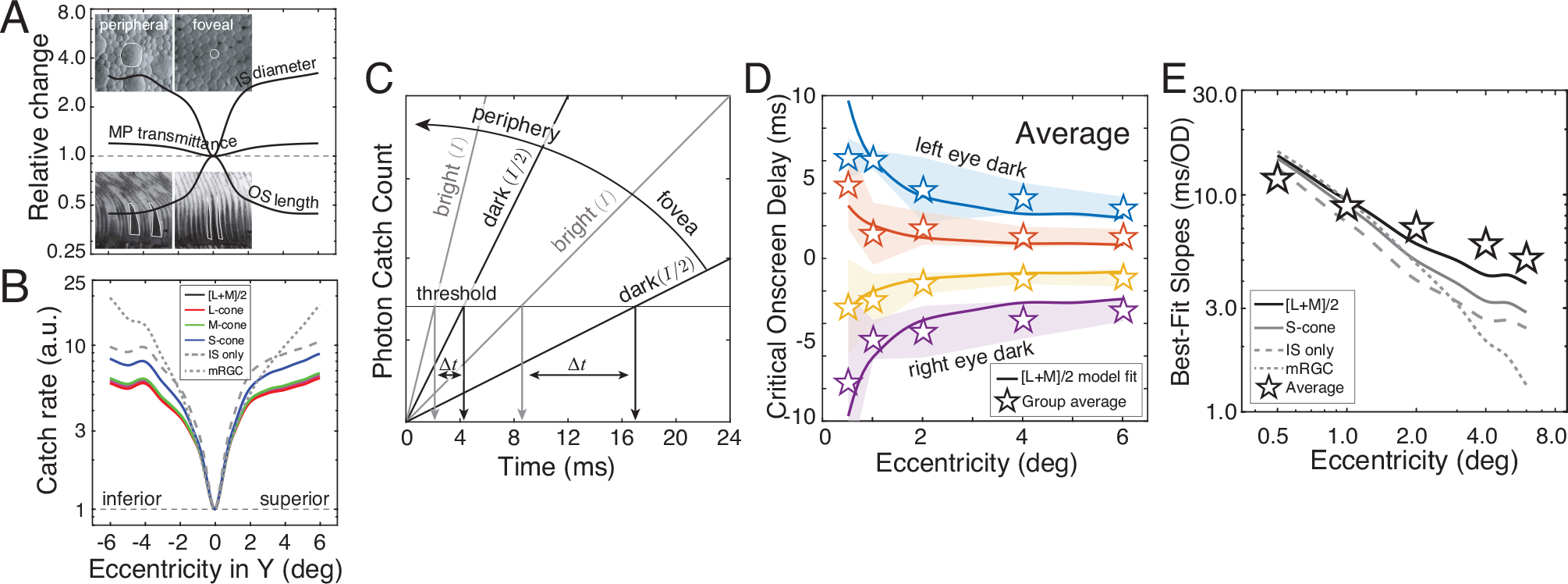
Retinal physiology-based model predictions. **A** Changes in retinal physiology with eccentricity, expressed as proportional change relative to the fovea. Cone inner-segment (IS) diameters increase (blue), cone outer-segment (OS) lengths decrease (red), and macular pigment transmittance (yellow) increases with eccentricity. Insets (adapted from Curcio et al., 1990) show peripheral and foveal cone IS diameters (top) and cone OS lengths (bottom), with individual examples outlined in white. **B** Photon catch rate, expressed as proportional change relative to the fovea, based on differences shown in A, for the L-, M-, and S-cones. The cone-specific catch rates differ because of how the cone spectral sensitivities interact with the spectral transmittance of the macular pigment, which passes less light in the short-than in the long-wavelength portion of the spectrum. Also shown are catch rates for the inner-segments only and for the midget ganglion cells (mRGCs). **C** Visualization showing how interocular delays (Δ*t*) are predicted from catch rates (slopes) for a 2-fold difference in light intensity (*I*) between the eyes. Higher catch rates in the periphery (see B) predict smaller interocular delays than in the fovea. **D** Model fit (solid curves) to group averaged data (stars) for all eccentricities and luminance differences. The model uses the mean catch rate of L- and M-cone catch rate (i.e. [L+M]/2). A single free-parameter—the threshold number of photons—was used to fit the data across all conditions. Changing the threshold value multiplicatively scales the model predictions symmetrically around zero delay. **E** Model fits (solid curves) and group averaged data replotted as in Fig. 2F. The [L+M]/2 model fit is the solid curve. Best-fit model predictions from S-cones (blue curve), inner-segment diameters alone (dashed curve), and midget ganglion cells (dotted curve) are also shown.

If peripheral cone photoreceptors absorb more photons per unit time—that is, have a higher photon ‘catch rate’—than foveal cone photoreceptors, perhaps the decrease in delay with eccentricity is straightforwardly accounted for. In line with available data, we assume that photon catch rate scales proportionally to i) the area of the inner-segments (Curcio et al., 1990), ii) the length of the outer-segments (because longer outer-segments contain more opsin molecules and hence absorb a greater proportion of incident photons) (Curcio et al., 1990; Banks et al., 1991; Renner et al., 2004), and iii) the transmittance of the macular pigment (Putnam & Bland, 2014). The scale factors relating these factors to photon catch rate are taken from the literature (Cottaris et al., 2019), and were implemented via the Image Systems Engineering Toolbox for Biology software package (ISETBio, http://isetbio.org).

Figure 4B plots the catch-rates of the L-, M-, and S-cones, and midget retinal ganglion cells, all relative to the foveal catches as predicted by the combined effect of the factors mentioned above. The catch rates of L-, M-, and S-cones are calculated by multiplying the spectral sensitivities of each cone type by the spectral transmittance function of the macular pigment, scaling by the influence of the inner-segment diameter and outer-segment length, and then integrating over wavelength. For purposes of comparison, we also include the catch rates for inner-segment diameters alone, and for midget retinal ganglion cells (mRGCs), presuming they pool the photons caught by the photoreceptors projecting to them.^2^

To develop an intuition of how the model converts catch rates into a prediction of interocular delay in each condition of our experiment, it is useful to visualize how the time required to catch the threshold number of photons depends on the catch rate (Fig. 4C). For a given light-level, and a given light-adapted state, the times to response in the bright and dark eyes are given by

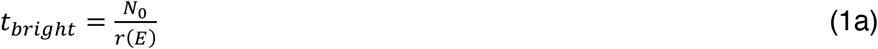

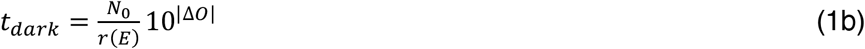

where *N*_0_ is the threshold number of photons that must be caught, *r*(*E*) is the catch rate as a function of eccentricity *E*, and 10^|Δ*O*|^ is the reciprocal of the transmittance *T* of the light from the monitor to the dark eye corresponding to the simulated optical density difference. The size of the model-predicted interocular delay

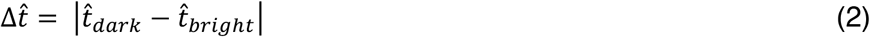

is given by the difference between the time to response in the two eyes. At a given eccentricity, the time to response is shorter in the bright eye than in the dark eye. And because catch rates increase with eccentricity, the model predicts shorter interocular delays at peripheral than at foveal visual locations (Fig. 4C).

Model predictions of the group averaged human data are shown in Fig. 4DE. The primary model uses the mean catch rates of the L- and M-cones (i.e. [L+M]/2) on the assumption that luminance sensitivity drives performance. A single free parameter, the threshold number of photons *N*_*0*_, is fit across all conditions to minimize mean absolute error (MAE). This free parameter multiplicatively scales the fits symmetrically about zero. So the data patterns that the model can account for are highly constrained. The human data is nevertheless well accounted for by the model (Fig. 4D).

Fig. 4E summarizes the patterns in the data as in Fig. 2F, for several additional models derived from the factors in Fig. 4AB. These models use the catch rates of the inner-segment diameters only, the S-cones, and the midget retinal ganglion cells (mRGCs). The [L+M]/2 model accounts for the group-averaged data—and for the individual subject data (see Supplementary Fig. S3)— better than the other models, and substantially so for all but the S-cone model.

Although the [L+M]/2 model is the best-fitting of the models, it predicts a slightly steeper decrease in delay than is observed in the psychophysical data. The [L+M]/2 model prediction is best-characterized by a power function with an exponent of -0.51, whereas the group-averaged psychophysical data is best-characterized by a power function with an exponent of -0.33 (see Fig. 2F). The retinal-model thus predicts a 2-fold decrease in delay with every 4-fold increase in eccentricity. (Recall that the psychophysical data indicates a 2-fold decrease in delay with every 8-fold increase in eccentricity). This mismatch may suggest that processing subsequent to the photoreceptors may partially compensate for spatio-temporal non-uniformities across the retina. Determining whether these compensatory processes are retinal (see below) or cortical, and understanding the principles on which they operate, is an interesting direction for future work.

The close match between the [L+M]/2 model predictions and the psychophysical results indicate that the catch rates of the cone photoreceptors can, to first approximation, account for temporal delays across a reasonable expanse of the central visual field. The match also provides additional support—and arguably more direct support than previous studies—for the consensus view that the temporal properties of retinal response drive the classic Pulfrich effect. However, this model leaves out several factors that could impact the predictions. First, it does not incorporate the effects of light-adaptation. This concern is mitigated by the interleaved experimental design which maintains the same light-adapted state in the visual system, on average, across conditions. Second, the model assumes that the temporal properties of photoreceptor response are determinative of the perceptual effects, even though it is the ganglion cells that carry signals from the eye to brain. This concern is mitigated by observations that ganglion cell kinematics, measured intracellularly at the cell body, are largely inherited from photoreceptor kinematics (Sinha et al., 2017). Third, the model does not include the fact that ganglion cell axonal conduction velocities differ as a function of retinal location. The concern that these differences might distort the model predictions is mitigated by the fact that signals needing to travel farther travel faster, at least partially compensating for the different distances that nerve impulses must travel to the optic nerve head before leaving the eye (Bucci et al., 2023). Nevertheless, including these factors in refined versions of the model may show that differences between the predictions and the psychophysical data are a consequence of having left these or other retinal factors out. Alternatively, it may be that post-retinal processes account for the discrepancy between the fits and the data. Still, the current model indicates that the reported psychophysical effects can be largely accounted for by major facets of the retinal physiology.

### Limitations and future directions

The current study is limited in several respects. Most obviously, it examined only how temporal delays caused by light-level differences are modulated by eccentricity. Images of natural scenes differ in many other ways (contrast, color, level of detail, blur) across space (see Fig. 1A). Future work should examine how eccentricity modulates the properties of temporal response to these other stimulus properties. Understanding how these factors combine to determine the kinematics of visual response to natural images is an ultimate goal, as these are the stimuli visual systems evolved to process (Burge, 2020; Burge & Geisler, 2011; 2012; 2014; 2015). Another limitation is that the radially-symmetric stimuli used in the experiments did not allow us to infer, for example, whether superior or inferior visual-field locations drive the effects. Appropriate modification of the experimental paradigm could allow more focal probing of temporal processing at specific visual field locations (e.g. x=-6º, y=+2º), which would facilitate comparison to more spatially-specific physiological results. Also, developing new paradigms—or modifying old ones (e.g. Prestrude & Baker, 1968)—that enable precise assessment of processing delays with monocularly-presented stimuli would increase the ease with which such measurements can be made and so increase how widely this topic is studied. Finally, updating the physiology-based model(s) to include response kinematics (e.g. impulse response functions) would be an ambitious, but potentially, useful undertaking.

Vision science has obtained a detailed understanding of what stimulus features are processed and signaled by stimulus sensors. Our understanding is much more limited of when those signals are processed and how asynchronies between them are resolved. The general problem of how to integrate multiple temporally-staggered signals is faced not just by the biological system, but by many human engineered systems, so this important but understudied problem presents an opportunity for further research.

## Conclusion

In this manuscript we report a precise characterization of how visual processing speed changes with eccentricity in the central portion (i.e. ±6º) of the visual field. A retinal-physiology-based model provides a tight quantitative account of the psychophysical data, suggesting that the photoreceptors themselves are the primary determinant of eccentricity-dependent processing delays. Intriguingly, variations in photoreceptor dynamics across the retinal surface are widespread in nature, and occur in creatures ranging from humans to flies (Burton et al., 2001; Masland, 2017; Sinha et al., 2022). Successful compensation for—or nulling of—differences in processing speeds must occur to prevent perceptual inaccuracies. The computational challenges that the human visual system must overcome to support accurate vision are substantial. Similar computational challenges are likely faced by visual systems across the animal kingdom.

## Methods

### Participants

Two male and three female subjects between the ages of 19 and 44 participated in the experiment. One male and one female subjects were authors; the other subjects were naïve to the purposes of the experiment. Visual acuity was normal or corrected-to-normal (i.e. visual acuity of 20/20 or better) in both eyes of each subject. Stereo-acuity was also normal, as assessed with the Titmus stereo test (i.e. stereoacuity better than or equal to 30arcmin). The experimental protocols were approved by the Institutional Review Board at the University of Pennsylvania and were in compliance with the Declaration of Helsinki. All participants provided written informed consent.

### Apparatus

Stimuli were presented on a four-mirror haploscope. Left- and right-eye images were presented on two identical VPixx VIEWPixx monitors. Monitors were calibrated (i.e., the gamma functions were linearized) using custom software routines. The monitors had a size of 53.3×30.0cm, a spatial resolution of 1920×1080 pixels, a native refresh rate of 120Hz, and a maximum luminance of 105.9cd/m2. After light loss due to mirror reflections, the maximum luminance was 93.9cd/m2. The two monitors were controlled by the same AMD FirePro D500 graphics card with 3GB GDDR5 VRAM, ensuring that the left and right eye images were presented synchronously. Custom firmware was written so that each monitor was driven by a single color-channel; the red channel drove the left monitor and the green channel drove the right monitor. The single-channel drive to each monitor was then replicated in all three-color channels to enable gray scale presentation.

Subjects viewed stimuli on the monitors through mirror cubes with 2.5cm circular openings positioned one inter-ocular distance apart. The field of view was approximately ±10º. The haploscope mirrors were adjusted such that the vergence distance matched the distance travelled by the light from the monitors to the eyes. The light travelled 100cm to the eyes, a distance confirmed both by laser ruler measurements and by binocular visual comparisons with a real target at 100cm. At this distance, each pixel subtended 0.93arcmin. Stimulus presentation was controlled via the Psychophysics Toolbox-3. Anti-aliasing enabled sub-pixel resolution permitting accurate presentations of disparities as small as 15-20arcsec. The head was stabilized with a chin and forehead rest.

### Stimuli

The target stimulus was a set of eight, equally-spaced circular beads that moved on one of five annulus-shaped paths. Each path was centered on a central fixation point. The left- and right-eye onscreen x- and y-positions of an individual circular bead were given by

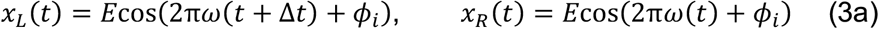

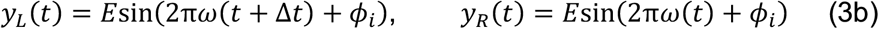

where *E* is the circular path’s radius (i.e. the retinal eccentricity), *ω* is the temporal frequency of the target movement cycles per second, *ϕ*_*i*_ is the starting phase, *t* is time in seconds, and Δ*t* is the onscreen delay between the left- and right-eye target images. The circular paths had radii of 0.5, 1.0, 2.0, 4.0, and 6.0º of visual angle. The temporal frequency always equalled ±1/*π* revolutions per second, so the beads moved either clockwise or anti-clockwise at constant speeds of 1.0, 2.0, 4.0, 8.0, or 12deg/sec, depending on the eccentricity. The starting phase of the first bead of eight was chosen randomly on the interval 0 to 2*π* (polar angle = 360deg). The other seven beads had equally spaced starting phases separated by *π*/4 (polar angle = 45deg). Trial duration was 0.25 seconds.

In the original experiment, the size of the beads increased in proportion to the eccentricity (i.e. radius) of the corresponding circular path (just as did the speeds). The circular beads had diameters of 10arcmin, 20arcmin, 40arcmin, 80arcmin, and 120arcmin at five eccentricities: 0.5º, 1.0º, 2.0º, 4.0º, and 6.0º.

In the size-control experiment (see below), the bead size was fixed to 20arcmin at all of the four probed eccentricities: 1.0º, 2.0º, 4.0º, and 6.0º. We did not use the smallest bead size from the main experiment (10arcmin) because it was hard to see at large eccentricities, and we did not probe the smallest eccentricity from the main experiment (0.5º) because 20 arcmin beads overlapped one another at the bead spacing used in the control experiment.

The onscreen interocular delay (negative or positive) and the direction of motion (clockwise or anti-clockwise) jointly determined whether onscreen stereo information specified whether the circular path was rotated ‘top back’ or ‘bottom back’ with respect to the screen (Fig. 1C). When the onscreen delay was negative, the left-eye stimulus was delayed onscreen relative to the right eye stimulus. When the onscreen delay was positive, the left eye stimulus was advanced onscreen. For clockwise motion, negative onscreen delays specified that the path was rotated ‘top back’ with respect to the screen and positive onscreen delays specified that the path was rotated ‘bottom back’ with respect to the screen. For anti-clockwise motion, the reverse was true.

Note that we did not temporally manipulate when left- and right-eye images were presented onscreen. Rather, we calculated the effective binocular disparity given the target velocity and the desired onscreen delay on each time step, appropriately shifted the spatial positions of the left- and right-eye images to create an equivalent onscreen disparity, and presented these disparate images synchronously on each monitor refresh.

The onscreen interocular delay Δ*t* can thus be equivalently expressed as an onscreen disparity at each point in time. The maximum onscreen horizontal disparity occurs at the top and the bottom of the circular path, where the horizontal component of the onscreen velocity is highest. At these locations (see Fig. 1C), the horizontal component of the velocity *ν*_*x*_ equals 2*Eπω*, the onscreen horizontal disparity in degrees of visual angle is given by

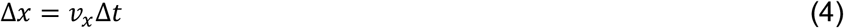

In this paper, uncrossed disparities are negative and crossed disparities are positive. Uncrossed and crossed disparities specify distances farther and closer than the screen, respectively.

Throughout the experiments, subjects fixated a white fixation dot in the center of the screen. The fixation dot was on top of a frontoparallel disk textured with 1/f noise. The diameter of the 1/f-textured disk equalled the diameter of circular path minus the diameter of the circular bead. The periphery of the display was also textured with 1/f noise, and had an opening with a diameter that equalled the diameter of the circular path plus the diameter of the circular bead. The space between the edge of the textured disk and the edge of the textured periphery thus had a width of twice the bead diameters and served as a ‘race track’ that the circular beads travelled along. Both the 1/f-textured disk and 1/f-textured screen periphery aided binocular fusion and served as stereoscopic references to the screen distance. Experiments were programmed in Matlab (Matlab 2017a; Mathworks, Inc.) and presented via PsychToolbox3 (Brainard, 1997).

### Procedure

Each set of moving circular beads was presented as part of a one-interval two-alternative forced-choice (2AFC) procedure. The task was to report, via a keypress, whether the circular path upon which the beads were traveling was rotated ‘top back’ or ‘bottom back’ with respect to the plane of the screen. Seven evenly spaced levels of onscreen interocular delay were presented with the method of constant stimuli. The particular levels of delay were set specifically for each subject in each condition based on pilot data.

The ‘top back’ and ‘bottom back’ responses were recorded as a function of motion direction (i.e. counter-clockwise and clockwise), onscreen interocular delay, interocular luminance difference, and retinal eccentricity. After appropriately collapsing across motion direction in each condition, we plotted the proportion of times subjects chose ‘top-back’ as a function of onscreen interocular delay for each eccentricity and luminance-difference condition (Fig. 2A). We refer to these data as the psychometric data functions (see below).

The experiment examined how eccentricity impacted the effect of luminance differences on processing speed as a function of retinal eccentricity. At each eccentricity, data was collected for each of four luminance differences. Either the left eye was delivered 75% or 50% less light than the right eye, or the right eye was delivered 50% or 75% less light than the left eye. Light losses of 75% and 50% correspond to (virtual) transmittances *T* of 0.25 and 0.50. The transmittance can be expressed in terms of an optical density *T* = 10^−*OD*^ where *OD* is the optical density of a ‘virtual’ neutral density filter. The virtual neutral density filter is implemented by reducing the luminance of the onscreen image by a scale factor equal to the transmittance *T*.

We express the luminance difference between the eyes in terms of the corresponding interocular difference in optical density

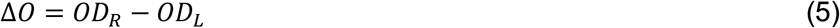

where *OD*_*L*_ and *OD*_*R*_ are the corresponding optical densities in the left and right eyes, respectively. The luminance differences used in the experiment correspond to optical density differences of Δ*O*=[-0.6, -0.3,+0.3,+0.6].

Data was collected at each eccentricity with all luminance difference conditions, all levels of interocular delay, and both motion directions interleaved in each of five blocks of 168 trials each (4 interocular image differences x 7 levels x 6 trials per level) for a total of 30 trials per level. Blocks probing different eccentricities were collected in pseudo-random counterbalanced order. Each block took approximately 3.5 minutes to complete. In total, at each eccentricity we collected a total 840 trials from each subject (210 trials per luminance difference x 4 luminance differences). In the main experiment, data was collected at five eccentricities (4200 trials per subject). In the size-control experiment (see below), data was collected at four eccentricities (3360 trials per subject).

### Data Analysis

The psychometric data functions were fit with a cumulative gaussian using maximum likelihood methods. The mean of the fitted psychometric function--that is, the onscreen delay that produced ‘top-back’ responses 50% of the time—is the estimated point of subjective equality (PSE). The point of subjective equality (PSE) is an estimate of the onscreen interocular delay necessary to make the circular beads appear to move within the screen plane, and hence should have equal size and opposite sign to the neural delays. Standard errors and/or confidence intervals were obtained for the PSE in each condition from 1000 bootstrapped datasets. The standard error of the PSE in a particular condition is given by the standard deviation of the bootstrapped distribution of PSE estimates.

We visualize the patterns in the PSEs—which are estimates of the critical onscreen delays—in two different ways: as a function of eccentricity for different luminance differences (Fig. 2BC) and as a function of interocular differences in optical density (Fig. 2DE). Recall that we use the optical density difference Δ*O* to quantify luminance differences between the eyes (see Eq. 5). At each eccentricity, the delays change linearly with luminance difference (Figs. 3DE).

The best-fit line at each eccentricity summarizes the effect of luminance difference Δ*O* on interocular delay at that eccentricity

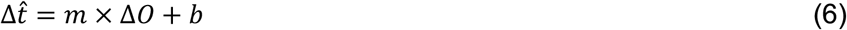

where *m* is the slope and *b* is the intercept of the best-fit line. We fit the data at each eccentricity with weighted linear regression, where the weights were set by the bootstrapped standard errors

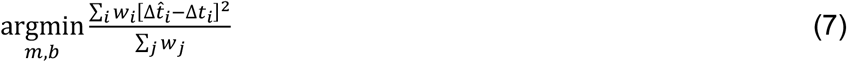

where *w* is the weight associated with a given observation (see Fig. 2E). Closed-form expressions exist for the best-fit parameters. The slopes of the best-fit regression lines are shown as a function of eccentricity in Fig. 2F for the grouped averaged data and for each individual subject, and in Fig. 3E for the group averaged data and various different retinal-physiology-based models.

To summarize how the slopes change with eccentricity, we fit a power function to the best-fit slopes across eccentricity with the objective of minimizing the squared distance between the fit and the data in log-space. This power function describes how the slopes change with eccentricity

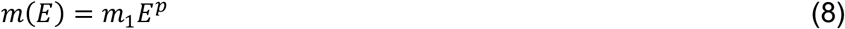

where *E* is the eccentricity, *p* is the power of the function, and *m*_1_ = *m*(*E* = 1) is the slope at the 1º of eccentricity. Combining Equations 6 and 8 yields a compact expression for interocular delays at all eccentricities and luminance differences

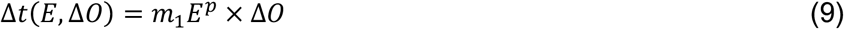

assuming that all y-intercepts (see Eq.5) are zero.

### Retinal Model

To examine whether properties of retinal physiology can account for the psychophysical data, we modeled how eccentricity-dependent changes in photoreceptor inner-segment diameters, outer-segment lengths, and macular pigment transmittance (Fig. 4A) affect photon catch rates of the cones (Fig. 4B), and then converted these photon catch rates into predictions of interocular delay.

The photon catch rate at an arbitrary eccentricity *E* relative to the foveal catch rate is given by

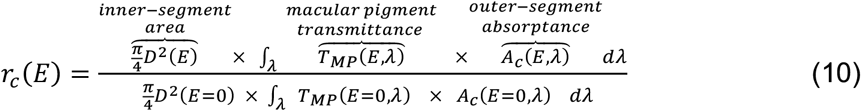

where *D* is the inner-segment diameter, *T*_*MP*_(*E, λ*) is the spectral transmittance of the macular pigment, *A*_*c*_(*E, λ*) is the spectral absorptance of each cone type *c* ∈ {*l, m, s*} that is due to combined effects of outer-segment length and the optical density and spectral sensitivity of the opsin molecules packed within it (see Cottaris et al., 2019). As in Cottaris et al. (2019), the foveal inner-segment diameter is taken to be 1.6 microns. Note that the numerator of Eq. 10 is the absolute catch rate at an arbitrary eccentricity and tha the denominator is the absolute catch rate at the fovea. The spectral absorptance of the outer-segment at each eccentricity and wavelength *λ* is given by

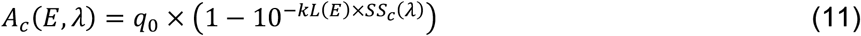

where *L*(*E*) is outer-segment length, *k* is the cone-specific density that converts length into axial optical density (Bowmaker & Dartnall, 1980), and *SS*_*c*_(*λ*) is the spectral sensitivity of each type of cone (Stockman & Sharpe, 2000; Stockman, Sharpe, Fach, 1999), and *q*_0_ is peak quantal efficiency. As in Cottaris et al. (2019), the cone specific density is taken to be 0.013 such that foveal outer-segment lengths of 38.5 microns yields a foveal axial optical density of 0.5. Also as in Cottaris et al. (2019), the peak quantal efficiency is taken to be 0.667 for all cone types. Note that, because the model relies on relative catch rates rather than absolute catch rates, and because the model assumes a constant uniform illumination, the specific values of some of the scale factors have no practical consequence for the model predictions.

### Retinal-model fitting

To convert the relative catch rates of the retinal model (see Eq. 10) to the pattern of critical onscreen delays across all conditions, we converted catch rate *r*(*E*) into interocular delay (see Fig. 4BC) as described in the main text (see Eqs. 1,2), and then fit a single free-parameter—the threshold number photons *N*_2_ required to elicit a photoreceptor response—to minimize the mean squared error (MSE) between the retinal-model-predicted and group averaged delays (Fig. 4D). Changing the threshold has the effect of multiplicatively scaling the model predictions of interocular delay symmetrically away from zero. The best-fits of several different models are shown in Fig. 4E. To ensure that the conclusions were not specific on the choice of error metric, we also fit the data to minimize the mean absolute error (MAE). Similar results were obtained.

## Acknowledgments

This work was supported by NIH grant R01-EY028571 from the National Eye Institute & Office of Behavioral and Social Science Research to J.B.. The authors thank Nicolas Cottaris for helpful discussion on cone photoreceptor physiology and for generous assistance with the ISETBio software package. The authors also thank Daniel Herrera and Josh Gold for comments on a draft version of the manuscript.

## Author Contributions

J.B. designed, directed, and performed research, contributed analytic tools, analyzed data, and wrote and edited the paper. C.M.D. designed and performed research, analyzed data, and wrote and edited the paper.

## Supplement

Delay discrimination thresholds are shown for all five human subjects (Fig. S1AB). For each observer, discrimination thresholds decrease with eccentricity. This result is consistent with an increase in temporal contrast sensitivity—which is well-known to improve in the peripheral visual field (Kelly, 1984). Also, like the critical onscreen delays, the delay discrimination thresholds are also reasonably well characterized by power functions, which plot as straight lines on log-log plots (Fig. S1B).

All five human subjects showed approximately the same pattern of critical onscreen delays in the main experiment and in the size-control experiment (Fig. S2A-C; also see Figs. 2,3). Four of the five subjects produced data that was clean enough to evaluate whether the effect of eccentricity in the original and size-control experiments was the same. For each of these four human subjects, the effect of eccentricity in each luminance difference condition is significantly correlated, and the best-fit regression lines to this data have slopes that were not significantly different 1.0. Data from the fifth subject is too noisy to draw firm conclusions (Fig. S2D). In general, the size-control experiment shows that our data does not support thinking that bead-size, rather than eccentricity, drove the pattern of visual processing delays.

**Figure S1.**
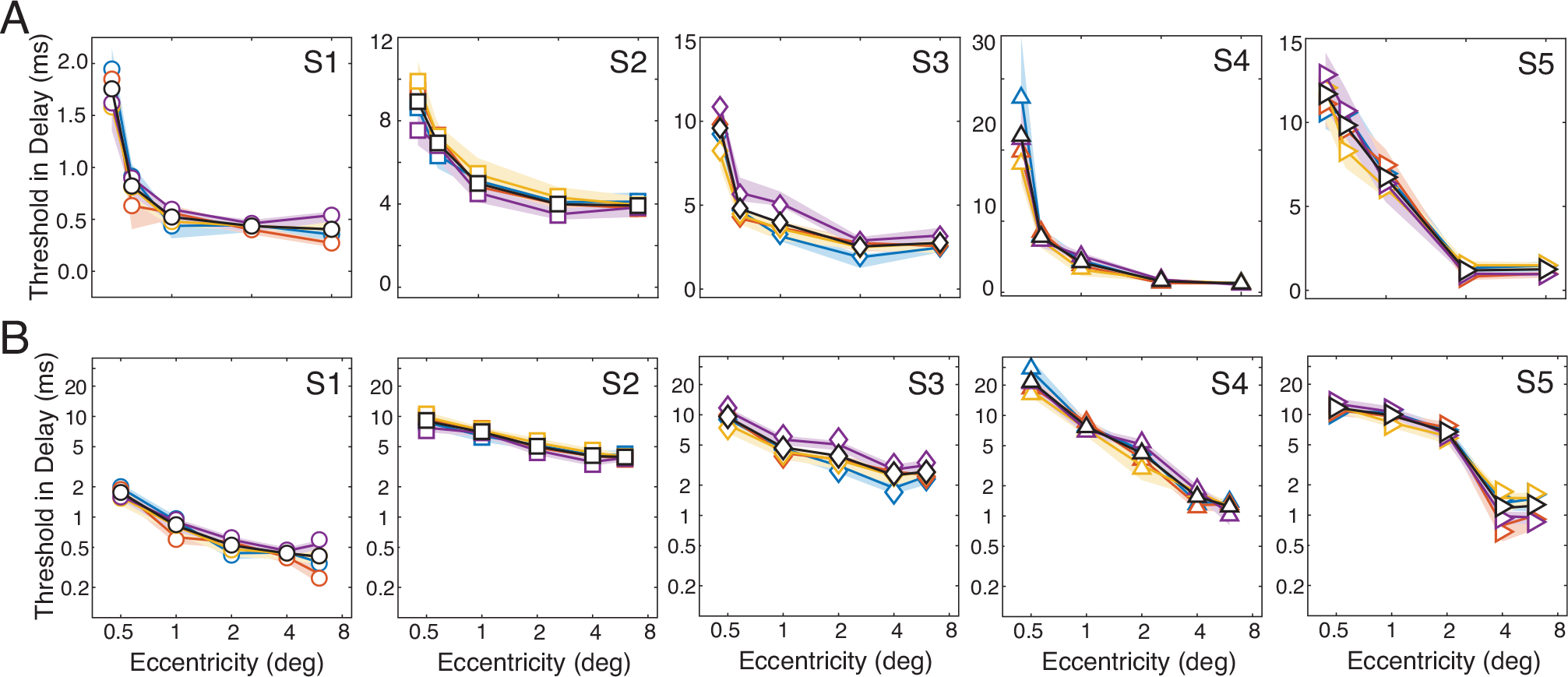
Individual subject thresholds. **A** Delay discrimination thresholds in milliseconds on linear-linear axes. Colors indicate luminance difference conditions (see Figs. 2-4) **B** Thresholds on log-log axes. Note that in A, the y-axis range different for each subjects whereas in B, it is the same for all subjects. The data is reasonably approximated by straight lines on the log-log plots indicating that— like the critical onscreen delays—a power function well-describes the change in threshold with eccentricity.

**Figure S2.**
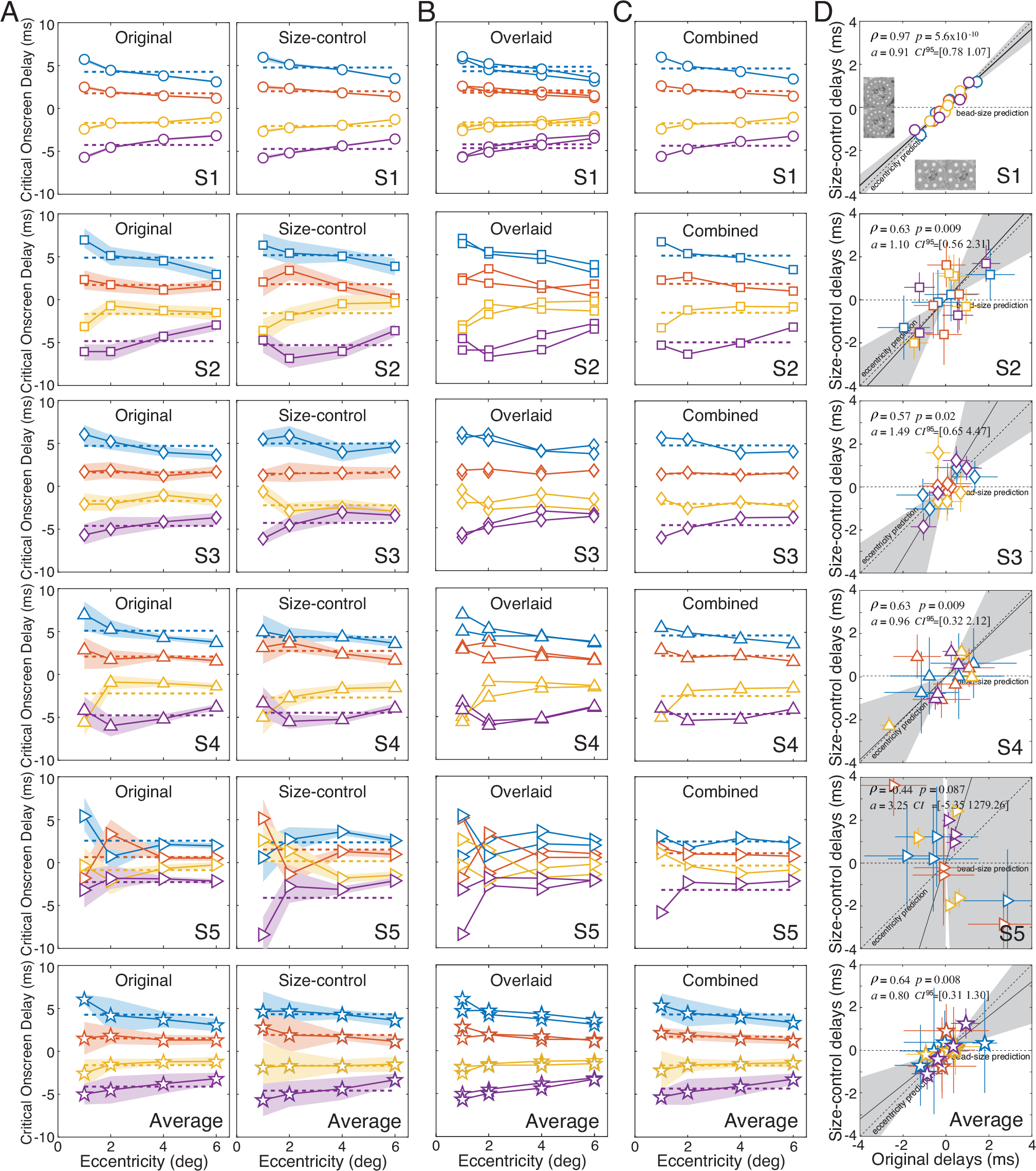
Individual subject, and average, data in original and size-control experiments. **A** Critical onscreen delays as a function of eccentricity at each luminance difference (colors) in both the original experiment (left) and the size-control experiment (right) for all five human subjects. Dashed lines indicate the mean delay at each luminance condition (colors). Each row contains the data for a different subject. **B** Overlaid data from A. **C** Critical onscreen delays averaged across the original and control experiments for each luminance difference and eccentricity. **D** Comparison of the effect of eccentricity in the original and size-control experiments, after regressing out the effect of luminance difference. The effect of eccentricity is significantly correlated in four of the five subjects, with best-fit slopes that are not significantly different from the unity line.

**Figure S3.**
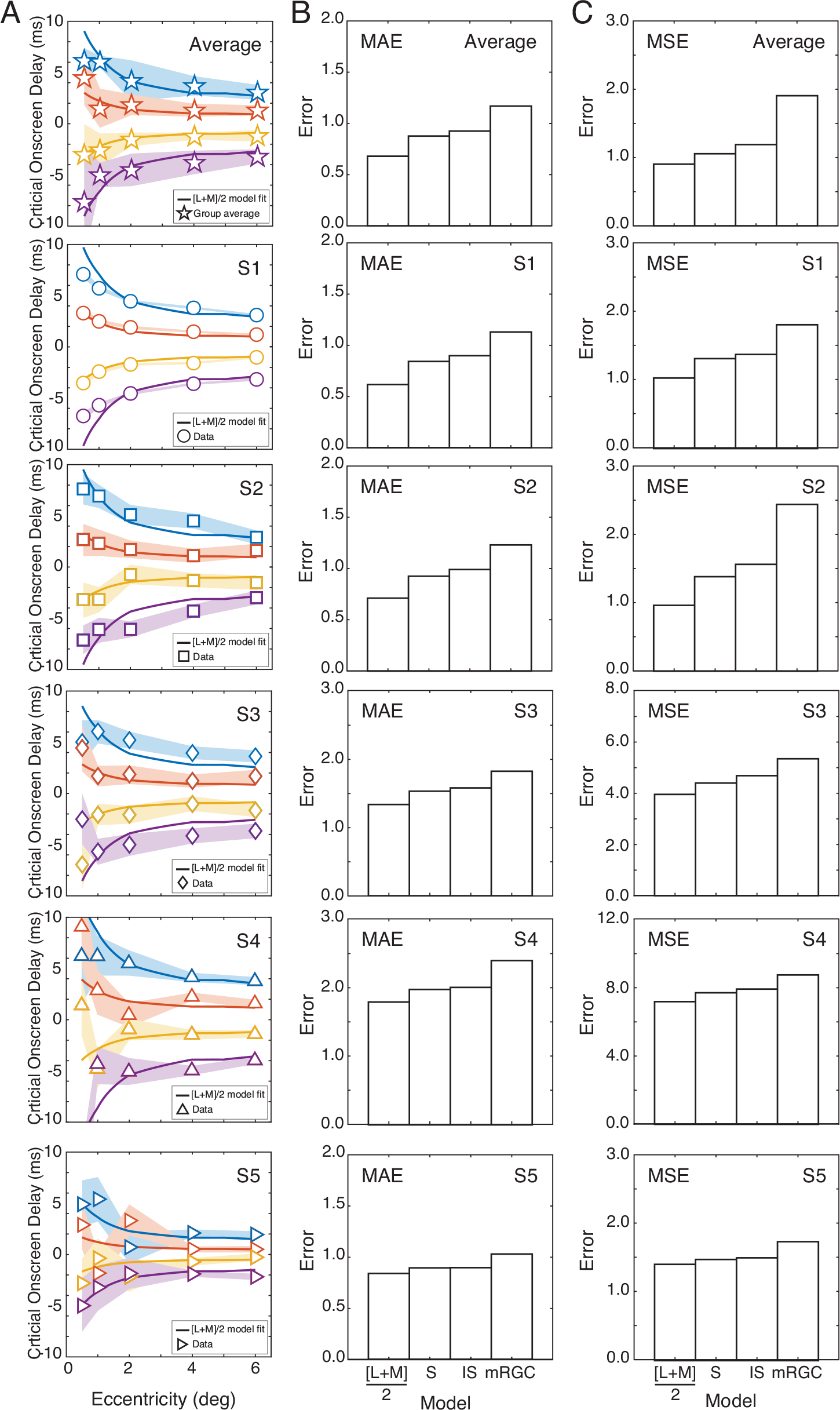
Model fits and model comparison. **A** Individual subject data and [L+M]/2 model-fits. Shaded regions indicate bootstrapped 68% confidence intervals (i.e. ∼+1 standard error). **B** Comparison of residual errors for the best-fit [L+M]/2 model, the S-cone model, the inner-segment diameter model, and the midget-retinal-ganglion-cell model using a minimum absolute error (MAE) objective. Note the fact that there are different y-axis scales for different subjects. **C** Same as B, but for a minimum mean squared error (MSE) objective.

## Footnotes

1 The current study-design does not rely on response-time measures, nor on comparisons of when in time foveal or peripheral targets are presented. Indeed, in the current study, temporal delays manifest as stereo-depth effects, so subjects are not required to make explicit estimates of any temporal aspect of the stimulus. This design feature is an advantage. People are often biased to overweigh foveally-presented information (Gloriani & Schütz, 2019). And response times—which are commonly used to assess processing latency— are often influenced by decision strategies, and inevitably include a motor contribution which may or may not depend on visual processing delays (Sternberg & Knoll, 1973). Determining the cause of response-time differences—visual, motor, or both—can thus be difficult. Stimulus-dependent changes in motor responses can sometimes be attributed to changes in front-end visual processing (Osborne et al., 2005; Lee et al., 2016; Chin & Burge, 2022; Burge & Cormack, under review). But such relationships must be demonstrated empirically.

2 The catch rate of the midget retinal ganglion cells (mRGCs) is obtained by scaling the mean catch rate of the L- and M-cones (i.e. [L+M]/2) by the number of cones in mRGC receptive-field-centers at each eccentricity (Watson, 2014). This computation implicitly assumes perfect spatio-temporal pooling of photon absorptions across the cones. The accuracy of this assumption is challenged by findings indicating that the dynamics of the mRGCs are largely inherited from the dynamics of the cone photoreceptors themselves (Sinha et al., 2017). The mRGC catch-rates, like the inner-segment only catch-rates, should thus be thought potentially useful benchmarks for comparison, rather than as realistic physiological possibilities.

